# Changes in ATP Sulfurylase activity in response to altered cyanobacteria growth conditions

**DOI:** 10.1101/2020.11.11.378158

**Authors:** Lucia Gastoldi, Lewis M. Ward, Mayuko Nakagawa, Mario Giordano, Shawn E. McGlynn

**Affiliations:** Laboratory of Algal and Plant Physiology, Department of Life and Environmental Sciences –(DISVA), Università Politecnica delle Marche (UNIVPM) - via Brecce Bianche, 60131 Ancona, Italy; Department of Earth and Planetary Sciences, Harvard University, Cambridge, Massachusetts, USA; Earth-Life Science Institute, Tokyo Institute of Technology - Ookayama, Tokyo, 152-8550, Japan

**Keywords:** ATPS, Sulfur, *Synechocystis*, *Synechococcus*, Proterozoic

## Abstract

Here we investigated variations in cell growth and ATP sulfurylase activity when two cyanobacterial strains – *Synechocystis* sp. PCC6803 and *Synechococcus* sp. WH7803 – were grown comparatively between conventional media and media with low ammonium, low sulfate and a controlled high CO_2_/low O_2_ atmosphere, which might resemble some Precambrian environments. In both organisms, a transition and adaptation to the reconstructed environmental media resulted in a decrease in ATPS specific activity. This decrease in activity appears to be decoupled from growth rate, suggesting the enzyme is not rate-limiting in S assimilation and raising questions about the role of ATPS redox regulation in cell physiology and thorughout history.

---

Sulfur is a universal and integral component of metabolism and biomass. Able to vary oxidation states from +6 to −2, and to populate the 3d orbitals, sulfur forms bonds with both carbon and iron, creating a bridge between the inorganic and organic in the cell (Beinert, 2000). With this electronic and molecular versatility, sulfur is involved in diverse and unique metabolisms across the tree of life (Dahl and Friedrich, 2008). At the same time, it is found in conserved co-factors and biomass components such as S-adenosyl methionine (Bridwell-Rabb et al., 2018; Marsh et al., 2010), coenzyme A (Strauss, 2010), and proteogenic cysteine residues, with their attendant in Fe-S clusters (Beinert et al., 1997; Gao, 2020; Rouault, 2019). With these proprieties, sulfur involving reactions likely had a prominent role from the origin of life onward (De Duve, 1991; Goldford et al., 2017; Russell et al., 1994; Wächtershäuser, 1990). Much later in evolution, sulfur availability in the oxidized form of sulfate may have influenced oceanic phytoplankton primary productivity (Giordano and Prioretti, 2016; Norici et al., 2005), and the radiation and evolution of this group (Ratti et al., 2011). Going further, sulfate variability in the ocean may also have recorded a linkage between animal evolution and the geochemical record of sulfate deposits (Canfield and Farquhar, 2009).

A considerable literature exists describing sulfur in the metabolism, ecology, and evolution of phytoplankton (Giordano et al., 2005, 2008; Kopriva and Rennenberg, 2004; Prioretti and Giordano, 2016; Takahashi et al., 2011). Some prominent areas of discussion are the sulfonium compound metabolism of DMSP/DMS (Giordano and Prioretti, 2016; Giordano et al., 2005; Ratti and Giordano, 2008; Takahashi et al., 2011), and protein Fe-S cluster biosynthesis (Cassier-Chauvat and Chauvat, 2014; Gao, 2020; Rina et al., 2000). In unicellular algae and cyanobacteria (as in many other organisms), S acquisition from the environment into biomass begins from sulfate (Giordano and Prioretti, 2016). Sulfate is kinetically inert and requires activation, which is then followed by reduction to biomass appropriate oxidation states (**Fig. 1**). **ATP Sulfurylase** (**ATPS** - EC 2.7.7.4) has the key role of SO_4_^2−^ activation at the beginning of S assimilation pathway, hydrolyzing ATP and producing a sulfate-ester (Giordano and Prioretti, 2016; Prioretti et al., 2014; Schmidt, 1972, 1988; Takahashi et al., 2011; Ullrich et al., 2001).

**Fig. 1.**
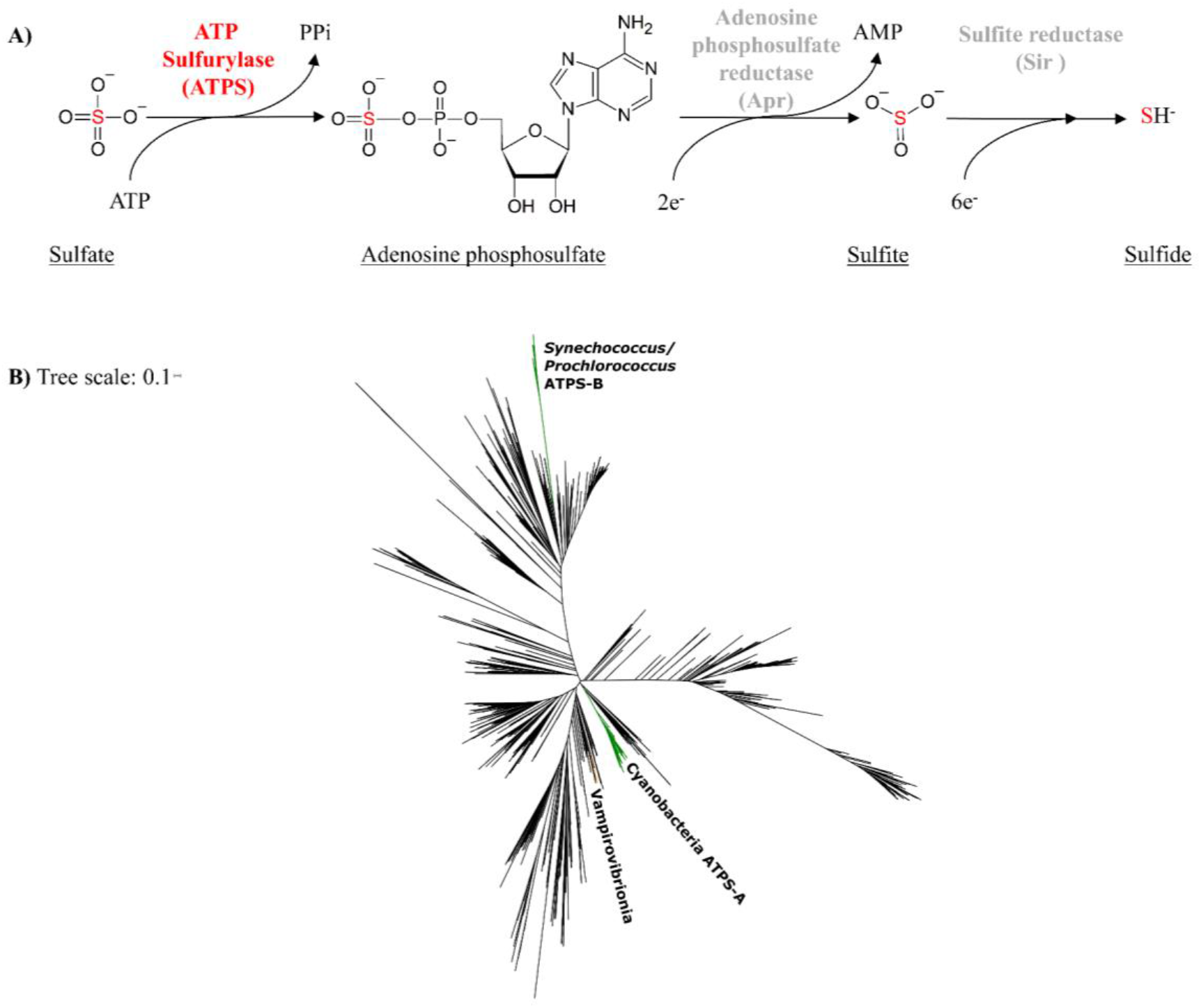
**A)** The role of ATP Sulfurylase in sulfate activation and the downstream steps leading to sulfide. The two step PAPS path can also reduce adenosine phosphosulfate to sulfite but is not shown. **B)** Phylogeny of ATP sulfurylase in bacteria and archaea. Homologs from the Vampiriovibrionia (an ancestral non-photosynthetic cyanobacteria class) are in orange, while the oxygenic cyanobacteria are labelled in green as the Synechococcus/Prochlorococcus clade with the ATPS-B isoform, and those cyanobacteria with the ATPS-A isoform. Remaining archaea and bacteria are colored in black. A more detailed tree can be found in the <u>Supplemental Materials</u>

In plants, S assimilation is regulated at different levels in response to growth conditions and in accordance with C and N metabolisms (Khan et al., 2010; Kopriva and Rennenberg, 2004; Koprivova and Kopriva, 2014; Martin et al., 2005; Takahashi et al., 2011; Vauclare et al., 2002). In both algae and cyanobacteria, diverse regulatory mechanisms exist (Schmidt, 1988; Takahashi et al., 2011) and have been investigated (Kopriva et al., 2002; MacRae et al., 2001; Patron et al., 2008; Prioretti et al., 2014). At the stage of sulfate activation, redox regulation of ATPS enzyme activity was postulated and subsequently confirmed (Prioretti et al., 2014, 2016). Within cyanobacteria, the redox regulated isoform, **ATPS–B**, contains 5 conserved cysteine residues (Prioretti et al., 2014), whereas **ATPS–A**, with only 4 conserved cysteine residues (Prioretti et al., 2014), appears to lack the critical regulatory S-residue organization thought to allow disulfide bridge formation and the modulation of enzyme activity.

It has been hypothesized that freshwater and marine cyanobacteria species are characterized by different evolutionary histories (Sánchez-Baracaldo, 2015; Sánchez-Baracaldo et al., 2017). Consequently, the different environments in which they evolved (perhaps including less or more oxidizing) could have influenced the presence/absence of redox regulation in specific proteins. The phylogenetic distribution pattern of ATPS homologs (**Fig. 1B** – see <u>Supplemental Material</u> for the explanation of how the tree was constructed) is suggestive this. The tree shows how the marine photosynthetic *Synnecococcus*/*Prochloroccoccus* (Syn/Pro) clade are well separated from other photosynthetic cyanobacteria (including the freshwater group). Notably, the Vampiriovibrionia class, since it represents an ancient and non-photosynthetic cyanobacteria taxon (Soo et al., 2019), was expected to be on a different branch and indeed it is possible to recognize as such (**Fig. 1B**). Within freshwater cyanobacteria species (and those marine species not enclosed in the Syn/Pro clade) the ATPS–A isoform without redox regulation is found, while, in the more derived Syn/Pro clade (which constitutes the picocyanobacteria plankton) the ATPS–B isoform with redox-regulation is found (Giordano and Prioretti, 2016; Prioretti et al., 2014, 2016). This pattern of distribution also corresponds to ribulose 1,5-bisphosphate carboxylase/oxygenase (RubisCO), and carboxysomes (Badger and Price, 2003; Rae et al., 2013): cyanobacteria with ATPS–A match the freshwater and brackish β-cyanobacteria which possess RubisCO-1B and β-carboxysomes; while those with ATPS–B coincide with the marine α-cyanobacteria that have RubisCO-1A and α-carboxysomes (Prioretti et al., 2016).

Protein phylogenies can be linked with gene transfer events during evolution, and several studies indeed confirmed the importance of horizontal gene transfer (HGT) during the evolution of the oxygenic photosynthesis in the cyanobacteria taxon (Fischer et al., 2016 and references in it). It has been observed for example that the Syn/Pro clade seems to have received a large number of genes via HGT from Proteobacteria: several genes involved in the formation of the α–carboxysome have been transfer from this group to Cyanobacteria along with bacteriochlorophylls synthesis genes (Bryant et al., 2012; Ward and Shih, 2020). The ATPS phylogeny (where Proteobacteria are more distributed across the tree; <u>Supplemental Materials</u>) is consistent with this protein having been part of this exchange (**Fig. 1B**), with the ATPS–B sequences nested within a clade primarily made up of proteobacterial sequences and very distant from the ATPS–A clade made up of other cyanobacteria. Moreover, a small number of Vampiriovibrionia species having a different version of ATPS protein is consistent with them acquiring it via HGT after their divergence from oxygenic Cyanobacteria, similar to how they’ve acquired those proteins involved in the aerobic respiration (Soo et al., 2017, 2019).

To further our understandings of the ATPS protein and its regulation in cyanobacteria, we grew two cyanobacteria species: the freshwater *Synechocystis* sp. PCC6803 (referred to simply as *Synechocystis* from now on) with non-redox regulated ATPS–A and, and the marine *Synechococcus* sp. WH7803 (referred to as *Synechococcus* from now on) with redox-regulated ATPS–B in multiple growth conditions and measured the resulting enzyme activity with a crude cell extract assay. The experiments allowed a comparison of enzyme activity levels between the two isoform types when expressed in cells exposed (and adapted) to the modern environment and a possible Precambrian condition. In particular, we considered the Proterozoic Eon, which lasted from 2.5Gyr to 0.6/0.5Gyr ago (refer to the following for more description: Fischer, 2008; Knoll and Nowak, 2017; Lyons et al., 2014; Rasmussen et al., 2008) and saw the oxygenation of Earth’s atmosphere (Lyons et al., 2014). During this period is where most evolutionary theories place the differentiation and radiation of the cyanobacteria taxon (Sánchez-Baracaldo, 2015; Schirrmeister et al., 2016; Shih et al., 2017 - despite most of the extant diversity of this taxon having been accumulated during the Phanerozoic Eon - Louca et al., 2018).

For each specie, 3 experimental conditions (with 3 biological replicates each) were set up in a growth chamber (12h light/12h dark cycle, temperature 20°C and light with a white LED lamp at 50μmol photon/m^2^s of irradiance). The cells were grown as semi-continuous cultures with the dilution volume based on growth rate data (Gastoldi et al in prep.) The three conditions analysed were: **Standard Condition** (**ST – Table S1**), the **Possible Proterozoic Condition** (**PPr – Table S2**) and the **Transitional Condition** (**TR – Table S3**). <u>AMCONA medium</u> (Fanesi et al., 2014) was used for the marine *Synechococcus* specie while the <u>BG11 medium</u> (Stanier et al., 1971) was used for the freshwater *Synechocystis*. All the liquid cultures were bubbled continuously with the atmosphere corresponding to the specific condition and experiments were performed after an adaptation period (2/3 months). The PPr condition had a higher CO_2_/O_2_ ratio than today (10’000ppm of CO_2_ ensured by a controlled gas system) while the TR and PPr growth media had a decrease in sulfate, a switch from nitrate to ammonium, and, in the case of the *Synechococcus*, lowered Fe concentrations compared to the ST. TR and PPr were identical except for the gas composition, with TR using air and PPr bubbled with a CO_2_ mix (details available in <u>Supplemental Materials</u>). These alterations might capture some historical variability during the Proterozoic Eon, but this can and should be debated.

Growth curves were determined as explained in other works (Gastoldi et al.; in preparation). ATPS activity was measured using crude cell extracts (refer to <u>Supplemental Materials</u> for a detailed procedure - which folllowed Giordano et al., 2000): the activity was observed spectrophotometrically at 25°C for 15 minutes and the linear phase of the assay was considered for the data analyses, as in previous works (Burnell, 1984; Giordano et al., 2000; Prioretti et al., 2016). The specific activity of ATPS (expressed in *μmol/min · ml^−1^*) in the crude extract was then normalized to the concentration of the protein (expressed in *mg/ml*) of the extract itself which was determined through the Lowry/Peterson technique (Lowry et al., 1951; Peterson, 1977).

For both organisms, the specific activity of the ATPS enzyme in the cells was very different between conditions. In *Synechocystis*, the mean activity value was 1459nmol/min mg^−1^ in the ST condition, higher than in the TR condition (164nmol/min mg^−1^ – ANOVA, *p-value* = 0.000753, Tukey as post doc, n=9, **Fig. 2**), where S, and Fe were lowered in concentration compared to the standard media and N changed from nitrate to ammonium. The TR condition showed a lower value than also PPr condition, where the value was 634nmol/min mg^−1^, though the difference between TR and PPr was less significant judged by a Tukey test (used as post hoc) which gave a higher *p-value* (*p-value* = 0.071).

**Fig. 2.**
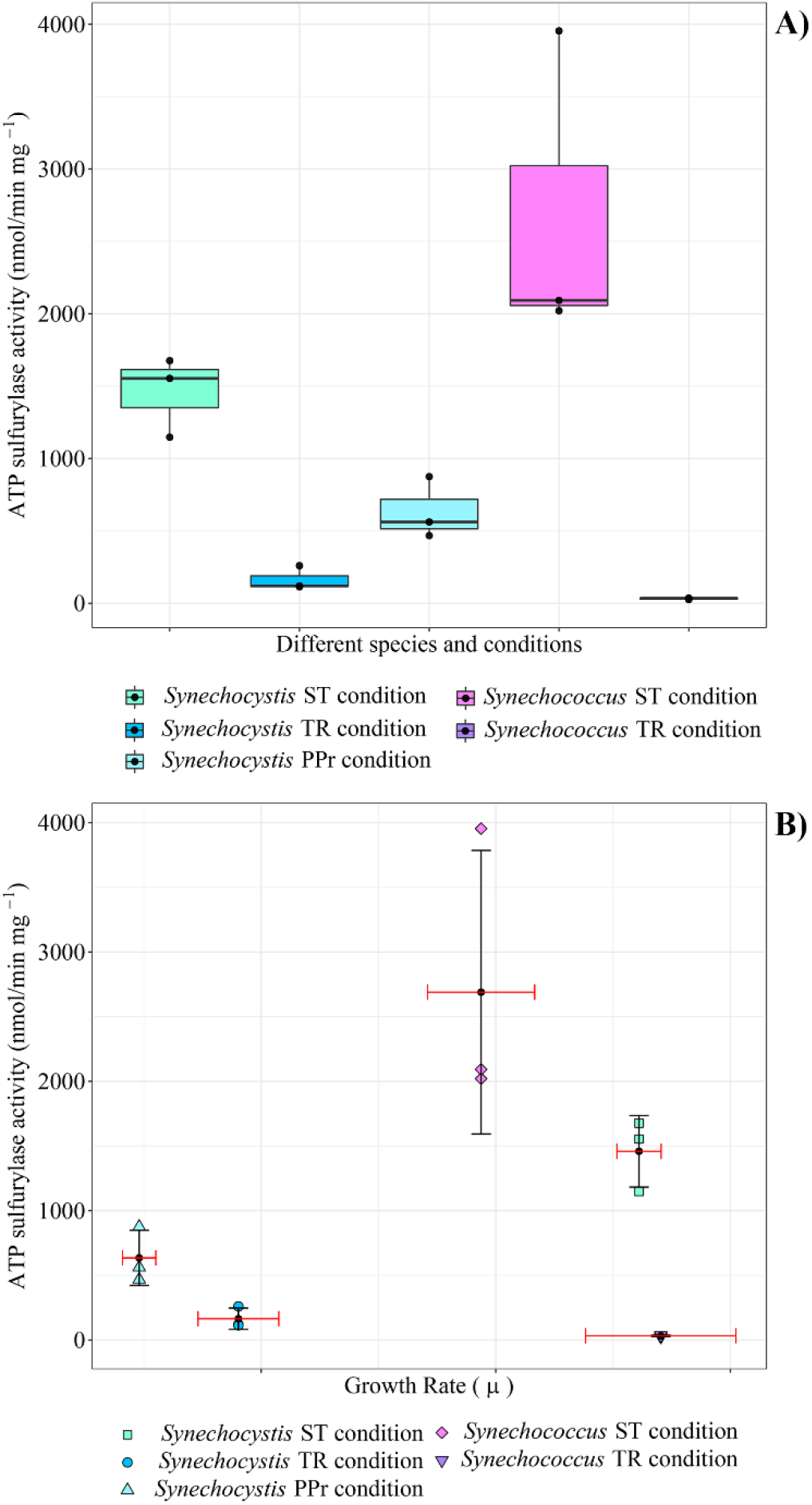
ATP sulfurylase activity. **A)** Each box represents the interquartile range of a specific condition and the black dots are the biological replicates for each one: the black line is the median and represents the second quartile Q2. Above the line is the upper quartile Q3 while below the line is the lower quartile Q1. The lines that come out from each box represent the minimum (lower) and the maximum (higher) value in the data. **B)** Each point represents a biological replicate; each condition is characterized with different shape and color. The black dot in each set of values is the mean of both the ATPS activity and the Growth Rate for the specific dataset. For each mean a Standard Deviation was also added, black SD for the ATPS activity value and red SD for the Grow Rate value. Single values used for this figure can be found in **Table S4** in the <u>Supplemental Material</u>

In *Synechococcus,* the average value was 2689nmol/min mg^−1^ in the ST condition while in the TR condition was only 32nmol/min mg^−1^ (t-Test *p-value* = 0.0137, n=6). The experiment was not possible for the seawater PPr condition since the *Synechococcus* specie was not able to survive in that condition despite several attempts for its adaptation.

In both *Synechocystis* and *Synechococcus*, the difference between the ATP sulfurylase activity in the ST and the TR conditions points out that media components other than supplied gas can have strong effects on the enzyme activity and growth, since the atmosphere was the same in both conditions. It may be that, in these organisms, a limitation in more than one nutrient (in our work we lowered sulfate and switched from nitrate to ammonium between ST and TR conditions), could decrease ATPS expression. This result is contrary to previous results where sulfate alone was varied and the activity increased in *Synechococcus* sp. WH7803 (Prioretti and Giordano, 2016). Further work is needed to address the factors that regulate ATPS, but we can now see that both nutrients supplied as dissolved salts can strongly affect the enzyme specific activity.

Plotting enzyme activity vs. growth rate for both the organisms, a trend between growth rate and ATPS activity was not observed, suggesting that the enzyme activity itself is not limiting growth in these conditions (**Fig. 2B**). In the marine *Synechococcus*, the growth rate in the TR condition was higher than the ST condition, but the ATPS enzyme activity was much lower, despite being in a medium with lower sulfate. In the case of *Synechocystis,* which could grow in the PPr condition and does not have a redox regulated ATPS (Prioretti et al., 2016), a higher value of ATPS activity was found in the PPr condition than in the TR condition, despite a lower growth rate value in the PPr condition (**Fig. 2B**). One hypothesis stemming from this observation is that the lower O_2_ available, which occurred in the PPr environment, promotes the activity of the enzyme or its expression. Although still preliminary, more data of this type – together with sulfur quota data which can be related with growth rates for some algae (Prioretti and Giordano, 2016) – will aid in assessing the relationships between ATPS enzyme activity, growth rate, and sulfur cell content. It is curious that in *Synechococcus,* while the enzyme activity decreased the S content of the cells increased (Gastoldi et al.; in prep). This could again imply that the enzyme is not limiting in the uptake of S. Together, our results pose questions as to why the enzyme is redox regulated in some organisms, and also why the activity varies so greatly between conditions.

We analyzed the phylogenetic distribution, and regulation of enzyme activity at the first step in ATP and electron requiring sulfate assimilation. By studying two organisms – one with redox regulation at the ATPS step and one without, our motivation was to gain insight into intra-cell redox responses in relationship to environmental redox. Our results highlight that cyanobacterial lineages display unique phenotypes in these conditions. This variability adds richness – and some complication – to theories about cyanobacteria evolution and adaptation to the oxygenation of the planet, as other works started recently to point out (Herrmann et al., 2020; Uchiyama et al., 2020). A long term goal is to investigate the possible linkages between metabolic regulation and Earth history.

## ACKNOWLEDGEMENTS

We want to thank the ELSI Research Interaction Committee for supporting the work, and both Tanaka Harumi and Nagano Reiko, for supporting the research logistics.

## References

Badger, M.R., and Price, G.D. (2003). CO2 concentrating mechanisms in cyanobacteria: molecular components, their diversity and evolution. J Exp Bot 54, 609–622.

Beinert, H. (2000). Iron-sulfur proteins: ancient structures, still full of surprises. JBIC 5, 2–15.

Beinert, H., Holm, R.H., and Münck, E. (1997). Iron-Sulfur Clusters: Nature’s Modular, Multipurpose Structures. Science 277, 653–659.

Bridwell-Rabb, J., Grell, T.A.J., and Drennan, C.L. (2018). A Rich Man, Poor Man Story of S-Adenosylmethionine and Cobalamin Revisited. Annu. Rev. Biochem. 87, 555–584.

Bryant, D.A., Liu, Z., Li, T., Zhao, F., Costas, A.M.G., Klatt, C.G., Ward, D.M., Frigaard, N.-U., and Overmann, J. (2012). Comparative and Functional Genomics of Anoxygenic Green Bacteria from the Taxa Chlorobi, Chloroflexi, and Acidobacteria. In Functional Genomics and Evolution of Photosynthetic Systems, R. Burnap, and W. Vermaas, eds. (Dordrecht: Springer Netherlands), pp. 47–102.

Burnell, J.N. (1984). Sulfate Assimilation in C4 Plants: Intercellular and Intracellular Location of ATP Sulfurylase, Cysteine Synthase, and Cystathionine β-Lyase in Maize Leaves. Plant Physiology 75, 873–875.

Canfield, D.E., and Farquhar, J. (2009). Animal evolution, bioturbation, and the sulfate concentration of the oceans. 5.

Cassier-Chauvat, C., and Chauvat, F. (2014). Function and Regulation of Ferredoxins in the Cyanobacterium, Synechocystis PCC6803: Recent Advances. Life 4, 666.

Dahl, C., and Friedrich, C.G. (2008). Microbial Sulfur Metabolism (Springer-Verlag).

De Duve (1991). Blueprint for a cell. RU Authors.

Fanesi, A., Raven, J.A., and Giordano, M. (2014). Growth rate affects the responses of the green alga Tetraselmis suecica to external perturbations. Plant Cell Environ 37, 512–519.

Fischer, W.W. (2008). Biogeochemistry: Life before the rise of oxygen. Nature 455, 1051–1052.

Fischer, W.W., Hemp, J., and Johnson, J.E. (2016). Evolution of Oxygenic Photosynthesis. Annual Review of Earth and Planetary Sciences 44, 647–683.

Gao, F. (2020). Iron–Sulfur Cluster Biogenesis and Iron Homeostasis in Cyanobacteria. Front Microbiol 11.

Giordano, M., and Prioretti, L. (2016). Sulphur and Algae: Metabolism, Ecology and Evolution. In The Physiology of Microalgae, (Springer, Cham), pp. 185–209.

Giordano, M., Pezzoni, V., and Hell, R. (2000). Strategies for the Allocation of Resources under Sulfur Limitation in the Green Alga Dunaliella salina. Plant Physiology 124, 857–864.

Giordano, M., Norici, A., and Hell, R. (2005). Sulfur and phytoplankton: acquisition, metabolism and impact on the environment. New Phytologist 166, 371–382.

Giordano, M., Norici, A., Ratti, S., and Raven, J.A. (2008). Role of Sulfur for Algae: Acquisition, Metabolism, Ecology and Evolution. In Sulfur Metabolism in Phototrophic Organisms, R. Hell, C. Dahl, D. Knaff, and T. Leustek, eds. (Springer Netherlands), pp. 397–415.

Goldford, J.E., Hartman, H., Smith, T.F., and Segrè, D. (2017). Remnants of an Ancient Metabolism without Phosphate. Cell 168, 1126–1134.e9.

Herrmann, H.A., Schwartz, J.-M., and Johnson, G.N. (2020). From empirical to theoretical models of light response curves - linking photosynthetic and metabolic acclimation. Photosynth Res 145, 5–14.

Khan, M.S., Haas, F.H., Samami, A.A., Gholami, A.M., Bauer, A., Fellenberg, K., Reichelt, M., Hänsch, R., Mendel, R.R., Meyer, A.J., et al. (2010). Sulfite Reductase Defines a Newly Discovered Bottleneck for Assimilatory Sulfate Reduction and Is Essential for Growth and Development in Arabidopsis thaliana. The Plant Cell 22, 1216–1231.

Knoll, A.H., and Nowak, M.A. (2017). The timetable of evolution. Sci Adv 3.

Kopriva, S., and Rennenberg, H. (2004). Control of sulphate assimilation and glutathione synthesis: interaction with N and C metabolism. J Exp Bot 55, 1831–1842.

Kopriva, S., Büchert, T., Fritz, G., Suter, M., Benda, R., Schünemann, V., Koprivova, A., Schürmann, P., Trautwein, A.X., Kroneck, P.M.H., et al. (2002). The Presence of an Iron-Sulfur Cluster in Adenosine 5′-Phosphosulfate Reductase Separates Organisms Utilizing Adenosine 5′-Phosphosulfate and Phosphoadenosine 5′-Phosphosulfate for Sulfate Assimilation. J. Biol. Chem. 277, 21786–21791.

Koprivova, A., and Kopriva, S. (2014). Molecular mechanisms of regulation of sulfate assimilation: first steps on a long road. Front. Plant Sci. 5.

Louca, S., Shih, P.M., Pennell, M.W., Fischer, W.W., Parfrey, L.W., and Doebeli, M. (2018). Bacterial diversification through geological time. Nature Ecology & Evolution 2, 1458–1467.

Lowry, O.H., Rosebrough, N.J., Farr, A.L., and Randall, R.J. (1951). Protein measurement with the Folin phenol reagent. J. Biol. Chem. 193, 265–275.

Lyons, T.W., Reinhard, C.T., and Planavsky, N.J. (2014). The rise of oxygen in Earth/’s early ocean and atmosphere. Nature 506, 307–315.

MacRae, I.J., Segel, I.H., and Fisher, A.J. (2001). Crystal structure of ATP sulfurylase from Penicillium chrysogenum: Insights into the allosteric regulation of sulfate assimilation. BIOCHEM.J. 40, 6795–6804.

Marsh, E.N.G., Patterson, D.P., and Li, L. (2010). Adenosyl Radical: Reagent and Catalyst in Enzyme Reactions. ChemBioChem 11, 604–621.

Martin, M.N., Tarczynski, M.C., Shen, B., and Leustek, T. (2005). The role of 5′-adenylylsulfate reductase in controlling sulfate reduction in plants. Photosynth Res 86, 309–323.

Norici, A., Hell, R., and Giordano, M. (2005). Sulfur and primary production in aquatic environments: an ecological perspective. Photosynth Res 86, 409–417.

Patron, N.J., Durnford, D.G., and Kopriva, S. (2008). Sulfate assimilation in eukaryotes: fusions, relocations and lateral transfers. BMC Evolutionary Biology 8, 39.

Peterson, G.L. (1977). A simplification of the protein assay method of Lowry et al. which is more generally applicable. Analytical Biochemistry 83, 346–356.

Prioretti, L., and Giordano, M. (2016). Direct and indirect influence of sulfur availability on phytoplankton evolutionary trajectories. J. Phycol. 52, 1094–1102.

Prioretti, L., Gontero, B., Hell, R., and Giordano, M. (2014). Diversity and regulation of ATP sulfurylase in photosynthetic organisms. Frontiers in Plant Science 5.

Prioretti, L., Lebrun, R., Gontero, B., and Giordano, M. (2016). Redox regulation of ATP sulfurylase in microalgae. Biochemical and Biophysical Research Communications 478, 1555–1562.

Rae, B.D., Long, B.M., Badger, M.R., and Price, G.D. (2013). Functions, Compositions, and Evolution of the Two Types of Carboxysomes: Polyhedral Microcompartments That Facilitate CO2 Fixation in Cyanobacteria and Some Proteobacteria. Microbiol. Mol. Biol. Rev. 77, 357–379.

Rasmussen, B., Fletcher, I.R., Brocks, J.J., and Kilburn, M.R. (2008). Reassessing the first appearance of eukaryotes and cyanobacteria. Nature 455, 1101–1104.

Ratti, S., and Giordano, M. (2008). Allocation of Sulfur to Sulfonium Compounds in Microalgae. In Sulfur Assimilation and Abiotic Stress in Plants, N.A. Khan, S. Singh, and S. Umar, eds. (Berlin, Heidelberg: Springer Berlin Heidelberg), pp. 317–333.

Ratti, S., Knoll, A.H., and Giordano, M. (2011). Did sulfate availability facilitate the evolutionary expansion of chlorophyll a+c phytoplankton in the oceans? Geobiology 9, 301–312.

Rina, M., Pozidis, C., Mavromatis, K., Tzanodaskalaki, M., Kokkinidis, M., and Bouriotis, V. (2000). Alkaline phosphatase from the Antarctic strain TAB5. European Journal of Biochemistry 267, 1230–1238.

Rouault, T.A. (2019). The indispensable role of mammalian iron sulfur proteins in function and regulation of multiple diverse metabolic pathways. Biometals 32, 343–353.

Russell, M.J., Daniel, R.M., Hall, A.J., and Sherringham, J.A. (1994). A hydrothermally precipitated catalytic iron sulphide membrane as a first step toward life. J Mol Evol 39, 231–243.

Sánchez-Baracaldo, P. (2015). Origin of marine planktonic cyanobacteria. Sci Rep 5.

Sánchez-Baracaldo, P., Raven, J.A., Pisani, D., and Knoll, A.H. (2017). Early photosynthetic eukaryotes inhabited low-salinity habitats. PNAS 114, E7737–E7745.

Schirrmeister, B.E., Sanchez-Baracaldo, P., and Wacey, D. (2016). Cyanobacterial evolution during the Precambrian. International Journal of Astrobiology 15, 187–204.

Schmidt, A. (1972). On the mechanism of photosynthetic sulfate reduction. Archiv. Mikrobiol. 84, 77–86.

Schmidt, A. (1988). [62] Sulfur metabolism in cyanobacteria. Methods in Enzymology 167, 572–583.

Shih, P.M., Hemp, J., Ward, L.M., Matzke, N.J., and Fischer, W.W. (2017). Crown group Oxyphotobacteria postdate the rise of oxygen. Geobiology 15, 19–29.

Soo, R.M., Hemp, J., Parks, D.H., Fischer, W.W., and Hugenholtz, P. (2017). On the origins of oxygenic photosynthesis and aerobic respiration in Cyanobacteria. Science 355, 1436–1440.

Soo, R.M., Hemp, J., and Hugenholtz, P. (2019). Evolution of photosynthesis and aerobic respiration in the cyanobacteria. Free Radical Biology and Medicine 140, 200–205.

Stanier, R.Y., Kunisawa, R., Mandel, M., and Cohen-Bazire, G. (1971). Purification and properties of unicellular blue-green algae (order Chroococcales). Bacteriol Rev 35, 171–205.

Strauss, E. (2010). 7.11 - Coenzyme A Biosynthesis and Enzymology. In Comprehensive Natural Products II, H.-W. (Ben) Liu, and L. Mander, eds. (Oxford: Elsevier), pp. 351–410.

Takahashi, H., Kopriva, S., Giordano, M., Saito, K., and Hell, R. (2011). Sulfur assimilation in photosynthetic organisms: molecular functions and regulations of transporters and assimilatory enzymes. Annu Rev Plant Biol 62, 157–184.

Uchiyama, J., Ito, Y., Matsuhashi, A., Ichikawa, Y., Sambe, M., Kitayama, S., Yoshino, Y., Moriyama, A., Kohga, H., Ogawa, S., et al. (2020). Characterization of Sll1558 in environmental stress tolerance of Synechocystis sp. PCC 6803. Photosynth Res.

Ullrich, T.C., Blaesse, M., and Huber, R. (2001). Crystal structure of ATP sulfurylase from Saccharomyces cerevisiae, a key enzyme in sulfate activation. EMBO J 20, 316–329.

Vauclare, P., Kopriva, S., Fell, D., Suter, M., Sticher, L., Ballmoos, P.V., Krähenbühl, U., Camp, R.O.D., and Brunold, C. (2002). Flux control of sulphate assimilation in Arabidopsis thaliana: adenosine 5′-phosphosulphate reductase is more susceptible than ATP sulphurylase to negative control by thiols. The Plant Journal 31, 729–740.

Wächtershäuser, G. (1990). Evolution of the first metabolic cycles. Proc Natl Acad Sci U S A 87, 200–204.

Ward, L.M., and Shih, P.M. (2020). Granick Revisited: Synthesizing Evolutionary and Ecological Evidence for the Late Origin of Bacteriochlorophyll via Ghost Lineages and Horizontal Gene Transfer. BioRxiv 2020.09.01.277905.

