## Supplemental Materials for "Changes in ATP Sulfurylase activity in response to altered cyanobacteria growth conditions"

| Condition | Modern Atmosphere | | Growth Media: Seawater AMCONA medium – *Synechococcus* – | | | Growth Media: Freshwater BG11 medium – *Synechocystis* – | | **Light** | **T°C** |
|---|---|---|---|---|---|---|---|---|---|
| | Gas | PAL (ppm) | Nutrients | [C] in the Medium | [C] in Modern Ocean | Nutrients | [C] in the Medium | | |
| Standard (ST) | $CO_2$ | ~407.8 | $Na_2SO_4$ | 25.0mM | 29mM | Nitrogen | $NaNO_3$ (17.65mM) | 50 µmol photon /m$_2$s | 20°C |
| | $O_2$ | ~209’460 | Nitrogen | $NaNO_3$ (549µM) | $NaNO_3$ (13.7µM) | $MgSO_4$ | 0.304mM | | |
| | $N_2$ | ~780’790 | $FeCl_3$ | 6.56µM | 2nM | Ammonium ferric citrate green | 0.6g/L stock (10ml stock/1L) | | |
| | | | $ZnSO_4$ | 254nM | 0.5nM | | | | |
| | | | $NaMoO_4$ | 105nM | 105nM | | | | |

***Table S 1*** *Description of the experimental condition defined as* <u>*Standard Condition (ST)*</u>*. The table shows the concentrations of fundamental elements, such as C, N, S, and Fe used for the AMCONA seawater medium (Fanesi et al., 2014) and BG11 freshwater medium (Stanier et al., 1971) flushing air with using air pump (KEDSUM-310 8W pump; Xiolan, China)*

| Condition | Possible Proterozoic Atmosphere | | Growth Media: Modified Seawater AMCONA medium – *Synechococcus* – | | Growth Media: Modified Freshwater BG11 medium – *Synechocystis* – | | **Light** | **T° C** |
|---|---|---|---|---|---|---|---|---|
| Possible Proterozoic (PPr) | Gas | ppm | Nutrients | [C] in PPr Medium | Nutrients | [C] in PPr Medium | 50 µmol photon /m$_2$s | 20° C |
| | | | $Na_2SO_4$ | 3mM | Nitrogen | $NH_4Cl_3$ (0.0035mM) | | |
| | $CO_2$ (20%) with 20ml/ min | 10’000ppm (~ 2’450% PAL) | Nitrogen | $NH_4Cl_3$ (100µM) | $MgSO_4$ | 0.035mM | | |
| | $O_2$ (in Air) with 5ml/ min | 20’000ppm (~ 10% PAL) | $FeCl_3$ | 200nM | Ammonium ferric citrate green | 0.6g/L stock 10ml stock/1L | | |
| | $N_2$ (100%) with 200ml/min | Base gas | $ZnSO_4$ | 0.0nM | | | | |
| | | | $NaMoO_4$ | 10.5nM | | | | |

***Table S 2*** *Description of the experimental condition defined as* <u>*Possible Proterozoic Condition (PPr)*</u>*. The table points out the modifications that were done to the AMCONA medium (Fanesi et al., 2014) and BG11 medium (Stanier et al., 1971) to mimic the Possible Proterozoic Environments*

| Condition | **Modern Atmosphere** | | **Growth Media** | | | | **Light** | **T°C** |
|---|---|---|---|---|---|---|---|---|
| | | | <u>Modified Seawater AMCONA medium</u> *– Synechococcus –* | | <u>Modified Freshwater BG11 medium</u> *– Synechocystis –* | | | |
| | Gas | PAL (ppm) | Nutrients | [C] in the PPr Medium | Nutrients | [C] in the Medium | | |
| Transitional (TR) | $CO_2$ | ~407.8 | $Na_2SO_4$ | 3mM | Nitrogen | $NH_4Cl_3$ (0.0035mM) | 50 µmol photon /$m_2$s | 20°C |
| | $O_2$ | ~209’460 | Nitrogen | $NH_4Cl_3$ (100µM) | $MgSO_4$ | 0.035mM | | |
| | $N_2$ | ~780’790 | $FeCl_3$ | 200nM | Ammonium ferric citrate green | 0.6g/L stock 10ml stock/1L | | |
| | | | $ZnSO_4$ | 0.0nM | | | | |
| | | | $NaMoO_4$ | 10.5nM | | | | |

***Table S 3*** *Description of the experimental condition defined as* <u>*Transitional Condition (TR)*</u>*. The table points out the modifications that were done to the AMCONA medium (Fanesi et al., 2014) and BG11 medium (Stanier et al., 1971) to mimic possible proterozoic Environments. This media has the same base liquid components as the PPr media in* ***Table S 2*** *but was used with a modern atmosphere*

| **Specie** | **Different condition** | **Activity** | | | **Growth Rate** | | |
|---|---|---|---|---|---|---|---|
| | | Biological replicates | Mean | SD | Biological replicates | Mean | SD |
| ***Synechocystis* sp. 6803** | Standard_1 | 1554 | | | 0.27 | | |
| | Standard_2 | 1147 | 1459 | 277 | 0.25 | 0.26 | 0.01 |
| | Standard_3 | 1676 | | | 0.26 | | |
| | Transitional_1 | 260 | | | 0.07 | | |
| | Transitional_2 | 120 | 164 | 82.6 | 0.10 | 0.09 | 0.02 |
| | Transitional_3 | 114 | | | 0.10 | | |
| | Possible Proterozoic_1 | 468 | | | 0.04 | | |
| | Possible Proterozoic_2 | 875 | 635 | 213 | 0.05 | 0.05 | 0.01 |
| | Possible Proterozoic_3 | 562 | | | 0.06 | | |
| ***Synechococcus* sp. 7803** | Standard_1 | 2021 | | | 0.17 | | |
| | Standard_2 | 2092 | 2689 | 1096 | 0.22 | 0.19 | 0.02 |
| | Standard_3 | 3954 | | | 0.19 | | |
| | Transitional_1 | 35.1 | 32.2 | 5.5 | 0.31 | 0.27 | 0.03 |
| | Transitional_2 | 25.8 | | | 0.25 | | |

| | | |
|---|---|---|
| Transitional_3 | 35.7 | 0.26 |

***Table S 4** Experimental data on which **Fig.2** is based on. SD indicates the Standard Deviation.*

**<u>ATPS assay</u>**

To start the extraction procedure, Extraction Buffer (containing 50mM of TRIS HCl pH8.1, 10mM of $MgCl_2$ and 1mM of EDTA - Burnell, 1984; Giordano et al., 2000; Prioretti et al., 2016) were added to the washed pellet and then, the cells were broken on ice using mortar and pestle. Subsequently, a solution of TritonX100 (0.1% v/v – to completely solubilize the proteins) and glycerol (10% v/v – to prevent protein degradation) was added to each sample. After 30 minutes of incubation on ice, the sample was slowly spun down at 4°C and the crude extract was collected and transferred to a new tube. 50µl from each tube were stored at -30°C to allow the quantification of the total amount of protein through the Lowry/Peterson procedure (Lowry et al., 1951; Peterson, 1977). The ATP sulfurylase activity was tested spectrophotometrically at 25°C for 15 minutes and only the linear phase of the assay was considered for the data analyses (Burnell, 1984; Giordano et al., 2000; Prioretti et al., 2016). The reaction mixture used for the enzyme test contained APS (1mM), PPi (as $Na_4P2O_7$, 1mM), $MgCl_2$ (5mM), glucose (5mM), NADP (300µM), hexokinase and glucose-6-P-dehydrogenase from baker's yeast (5units/ml, BIOCON JAPAN, LTD), and Tris HCl pH8.1 (50mM). During the analyses, the increase in NAPDH was detected at 340nm spectrophotometrically with the EnSpire Multilabel Reader (PerkinElmer).

**<u>Phylogenetics Methods</u>**

Phylogenetic analyses followed methods described previously (e.g. Ward and Shih, 2020) and summarized briefly here. Representative genomes from across the known diversity of bacteria and archaea were selected from the GTDB database (Chaumeil et al., 2020) dereplicated at the genus level. Genomes were downloaded from the NCBI WGS and Genbank databases. ATPS protein sequences were extracted using the tblastn function of BLAST+ (Camacho et al., 2009) and aligned with MAFFT (Katoh et al., 2009)and MUSCLE (Edgar, 2004). Trees were built with RAxML v.8.2.12 (Stamatakis, 2014)on the Cipres science gateway (Miller et al., 2010). Transfer bootstrap support values were determined with

BOOSTER (Lemoine et al., 2018). Visualization of trees was performed with the Interactive Tree of Life Viewer (Letunic and Bork, 2016). The tree can be observed in below.

**BIBLIOGRAPHY**


Burnell, J.N. (1984). Sulfate Assimilation in C4 Plants: Intercellular and Intracellular Location of ATP Sulfurylase, Cysteine Synthase, and Cystathionine β-Lyase in Maize Leaves. Plant Physiology *75*, 873–875.

Camacho, C., Coulouris, G., Avagyan, V., Ma, N., Papadopoulos, J., Bealer, K., and Madden, T.L. (2009). BLAST+: architecture and applications. BMC Bioinformatics *10*, 421.

Chaumeil, P.-A., Mussig, A.J., Hugenholtz, P., and Parks, D.H. (2020). GTDB-Tk: a toolkit to classify genomes with the Genome Taxonomy Database. Bioinformatics *36*, 1925–1927.

Edgar, R.C. (2004). MUSCLE: multiple sequence alignment with high accuracy and high throughput. Nucleic Acids Res *32*, 1792–1797.

Fanesi, A., Raven, J.A., and Giordano, M. (2014). Growth rate affects the responses of the green alga Tetraselmis suecica to external perturbations. Plant Cell Environ *37*, 512–519.

Giordano, M., Pezzoni, V., and Hell, R. (2000). Strategies for the Allocation of Resources under Sulfur Limitation in the Green Alga Dunaliella salina. Plant Physiology *124*, 857–864.

Katoh, K., Asimenos, G., and Toh, H. (2009). Multiple alignment of DNA sequences with MAFFT. Methods Mol Biol *537*, 39–64.

Lemoine, F., Domelevo Entfellner, J.-B., Wilkinson, E., Correia, D., Dávila Felipe, M., De Oliveira, T., and Gascuel, O. (2018). Renewing Felsenstein's phylogenetic bootstrap in the era of big data. Nature *556*, 452–456.

Letunic, I., and Bork, P. (2016). Interactive tree of life (iTOL) v3: an online tool for the display and annotation of phylogenetic and other trees. Nucleic Acids Res *44*, W242-245.

Lowry, O.H., Rosebrough, N.J., Farr, A.L., and Randall, R.J. (1951). Protein measurement with the Folin phenol reagent. J. Biol. Chem. *193*, 265–275.

Miller, M.A., Pfeiffer, W., and Schwartz, T. (2010). Creating the CIPRES Science Gateway for inference of large phylogenetic trees. In 2010 Gateway Computing Environments Workshop (GCE), pp. 1–8.

Peterson, G.L. (1977). A simplification of the protein assay method of Lowry et al. which is more generally applicable. Analytical Biochemistry *83*, 346–356.

Prioretti, L., Lebrun, R., Gontero, B., and Giordano, M. (2016). Redox regulation of ATP sulfurylase in microalgae. Biochemical and Biophysical Research Communications *478*, 1555–1562.

Ratti, S., Knoll, A.H., and Giordano, M. (2011). Did sulfate availability facilitate the evolutionary expansion of chlorophyll a+c phytoplankton in the oceans? Geobiology *9*, 301–312.

Stamatakis, A. (2014). RAxML version 8: a tool for phylogenetic analysis and post-analysis of large phylogenies. Bioinformatics *30*, 1312–1313.

Stanier, R.Y., Kunisawa, R., Mandel, M., and Cohen-Bazire, G. (1971). Purification and properties of unicellular blue-green algae (order Chroococcales). Bacteriol Rev *35*, 171–205.

Ward, L.M., and Shih, P.M. (2020). Granick Revisited: Synthesizing Evolutionary and Ecological Evidence for the Late Origin of Bacteriochlorophyll via Ghost Lineages and Horizontal Gene Transfer. BioRxiv 2020.09.01.277905.
